# Mind the instructions: reward cues are liked first, wanted later

**DOI:** 10.1101/2024.01.09.574803

**Authors:** Nicoleta Prutean, Luc Vermeylen, Nanne Kukkonen, S. Tabitha Steendam, Joshua O. Eayrs, Ruth M. Krebs, Jan R. Wiersema, Eliana Vassena, C. Nico Boehler, Wim Notebaert

## Abstract

Current theories propose that mental effort is invested only when the anticipated benefits, such as rewards, outweigh the associated costs, like task difficulty. Yet, it remains unclear whether this motivational and mitigating aspect of reward processing is reflected in the evaluation of reward/difficulty cues as such, and to what extent it depends on task experience. In a pre-registered experiment (N=84), we used the affect misattribution procedure (AMP) to gauge affective evaluations of nonword cues predicting reward and task difficulty levels. Contrary to previous studies, the AMP was administered at the outset, after cue instructions, and after the cues were used in a random dot motion (RDM) task. Compared to baseline, cues predicting a larger reward were evaluated more positively after RDM task experience, and most importantly, already after cue instructions, with no difference between the two phases. This evaluative effect manifested in increased performance after larger reward cues in the RDM task. Our results suggest that AMP effects may generally capture performance expectations which are independent of task experience. Importantly, these instructed expectations of reward and difficulty play a crucial role in the evaluation and subsequent investment of mental effort.

## 1. Introduction

Our modern daily lives consist of many mental challenges, ranging from minor tasks like remembering to pick up groceries to some bigger tasks like revising a scientific paper. This latter task demands considerable mental effort and is typically perceived as a negative experience. However, this perception can shift when reward is looming, as it happens, for example, when we are about to publish a paper that might be important for our future academic career (the reward). In this scenario, the potential reward is evaluated as positive and this counterbalances the initial negative evaluation of the effort involved.

While this scenario is familiar, extensive research in the fields of neuroeconomics (e.g., Westbrook et al., 2013), cognitive (e.g., Kool et al., 2010; Kukkonen et al., 2023) and computational neuroscience (Grahek et al., 2023; Shenhav et al., 2013, 2021; Silvestrini et al., 2023; Verguts et al., 2015) is currently focused on comprehending the precise mechanisms underlying how, why, and when we feel motivated to invest mental effort. Researchers concur that mental effort comes with an intrinsic cost, whether it be metabolic (Baumeister & Heatherton, 1996; Holroyd, 2015), computational (Shenhav et al.; Silvetti et al., 2018) or related to missed alternative opportunities (Kurzban et al., 2013). Nevertheless, people exert effort whenever potential benefits (e.g., rewards) outweigh the associated costs. This implies that effort investment is preceded by a preliminary evaluative phase, during which anticipated benefits are evaluated more positively, mitigating the challenges of investing mental effort.

According to the expected value of control theory (EVC, Shenhav et al., 2013; 2017), effort investment is the result of integrating information about costs and benefits from both external (e.g., cues, instructions, learned contingencies) and internal sources of information (e.g., motivational state). For example, when task difficulty and reward are cued on each trial, participants (obviously) perform worse on hard tasks, but they perform better on those trials when large rewards are available (Vassena et al., 2019, Experiment 1; see also Krebs et al., 2012; Schevernels et al., 2014). Moreover, when given the choice to perform tasks with different levels of difficulty and rewards, participants prefer easy tasks, but accept hard tasks more often if a larger reward is at stake (Vassena et al., Experiment 2; Westbrook & Braver, 2013). These results highlight the efficacy of incentives in stimulating motivation, intended as goal-directed behaviour aimed at minimising costs and maximising benefits (Berridge, 2004). However, Berridge & Robinson (2003) emphasize that there exists a separate and distinct (neural) process related to the affective experience tied to a reward (‘liking’), which is different from the motivation or drive induced by reward prospects (‘wanting’). For instance, whilst the dopaminergic mesolimbic system has been related to motivated behaviour (e.g., Berridge & Robinson, 1998; Krebs et al., 2012; Vassena et al., 2014), its activation or suppression does not influence the liking component (Peciña et al., 1997). In the same vein, Devine et al., 2023 have shown ‘liking’/‘disliking’ facial expressions (i.e., decreased/increased corrugator activity, respectively) in response to difficulty and reward cues and during task execution, but in the absence of any reward modulation of actual task performance. If trial-by-trial affective reactions to reward (‘liking’) are independent of task performance (‘wanting’), it remains to be seen whether affective evaluations leading to effort investment reflect broader expectations of reward and difficulty prior to task exposure.

In the current study, we used the affect misattribution procedure (AMP; Payne et al., 2005) to measure the affective value of reward and difficulty cues before and after experiencing their predictive value in a separate task. Similar to other paradigms (e.g., ‘affective priming’ in Fazio, 2001), the AMP allows estimation of the affective value of a stimulus. In this procedure, participants have to judge targets preceded by primes as pleasant/unpleasant. Crucially, the target is intended to be an ambiguous stimulus (e.g. a Chinese pictograph presented to non-Chinese-speaking participants), lacking any intrinsic value. On the contrary, the primes are thought to have an affective dimension, which becomes misattributed (Payne & Lundberg, 2014) to the contiguous target (AMP effect). Vermeylen et al. (2019, 2022) used the AMP to demonstrate that cues predicting (more effortful) task switches were evaluated as more negative compared to repetition cues after a task-switching paradigm (Monsell, 2003). Moreover, participants who were worse on switch trials also evaluated the switch primes more negatively, suggesting that the evaluation depended on participants’ experience during the task switching paradigm. Interestingly, Van Dessel et al. (2020) showed that mere cue instructions, without actual task experience, are sufficient to elicit a similar AMP effect of task switching cues. However, in their experiment, AMP was measured after participants had completed a short task switching block (although with different cues). Even though it is not possible to exclude any impact of experience on the effect, their experiment suggests that expectations based on solely instructions can have a role in the affective evaluation of anticipatory task cues. Our participants performed a random dot motion task (RDM; Rajananda et al., 2018), which allows subtle and individually tailored manipulations of task difficulty. In our task, participants responded to the motion direction of a cloud of dots moving coherently towards the left or right against a random dot motion background. Task difficulty was determined by the percentage of dots moving coherently, and it was adjusted individually through a staircase procedure to ensure a consistent level of performance across participants. Crucially, we manipulated reward prospect as well to measure their positive impact on mental effort evaluation and subsequent investment. Bidimensional nonword cues (e.g., ‘OXEYA’) prospecting task difficulty (easy or hard) and reward (small or large) were presented before each RDM trial. We then used the cues as primes to measure the evaluation (AMP) at the beginning of the experiment to account for any a-priori bias in pictograph evaluations (AMP baseline), after being instructed on the meaning of the cues, but in the absence of any task experience (AMP instruction), and after experiencing the predictive value of cues in the RDM task (AMP experience).

In the RDM task, we anticipated better performance when large rewards were prospected compared to small rewards (i.e., reward effect) and on easy trials compared to hard trials (i.e., difficulty effect). Moreover, we predicted a more pronounced impact of reward prospects on hard trials, with participants exhibiting their poorest performance when anticipating a small reward for a hard trial (i.e., interaction, as in Vassena et al., 2019). We hypothesized that the impact of reward prospects on performance (and potential interaction with difficulty) would also be reflected in their evaluations of those cues. Specifically, we expected more negative evaluations for cues prospecting small compared to large rewards (i.e., reward effect), for hard compared to easy trials (i.e., difficulty effect), and that even more so if for a small reward (i.e., interaction; AMP effects). In line with previous studies (Vermeylen et al., 2019; 2022), we anticipated AMP effects after experience with the cues in the RDM task (AMP experience vs AMP baseline), but, crucially, also based on solely cue instructions (AMP instructions vs AMP baseline).

The hypotheses and methods of the current study were preregistered at https://osf.io/nxdvh.

## 2. Methods

### 2.1. Participants

Our initial sample consisted in 100 participants recruited through SONA system at Ghent University (www.ugent.sona-systems.com)^1^ in exchange for a monetary compensation (13 euro as base payment + up to 7 euro reward based on performance). All participants were required to be between 18-35 years old, right-handed, have normal or corrected-to-normal vision, no history of psychiatric/neurological disorders, and no knowledge of Chinese characters. After applying our exclusion criteria (see Pre-processing paragraph below), 84 participants remained (of which 68 females, Mage = 22.5, SDage = 3.6).

### 2.2. Stimuli and procedure

Participants completed the AMP in three different phases: with meaningless nonword cues as primes (AMP baseline), after being instructed (cue instruction phase) on the reward/difficulty prediction of the nonword cues (AMP instructions), and after experiencing the predictive value of the cues in a RDM task (AMP experience). At the end of the experiment, they filled in some questionnaires measuring individual differences (see below).

In each affect misattribution procedure (Figure 1, left panel), 80 pseudo Chinese pictographs (randomly generated via http://generator.lorem-ipsum.info/_chinese) were used as targets, and were repeated twice across 4 different blocks (160 trials). Given that people tend to associate positive concepts with their dominant body side (Casasanto, 2009), participants were asked to evaluate the targets as rather unpleasant or pleasant by pressing the C or N keys respectively, with their left or dominant hand. Targets were preceded by non-word cues (“AFUBU”, “YOVIN”, “OXEYA” or “UPUSU”), which were previously rated as neutral by an independent sample (Mertens et al., 2018) and used in the same paradigm (Van Dessel et al., 2020; Vermeylen et al., 2019, 2022). Cues were randomly associated with pictographs and evenly presented (40 trials each). Each trial started with a fixation cross (500 ms), followed by the cue prime (400 ms), the target pictograph (until response or max 2000 ms) and an interval (500 ms). To ensure that the cues were processed, ∼ 10% of the trials were catch trials. On those trials, instead of evaluating the pictographs, participants had to select among two options the one corresponding to the just-presented cue identity (AMP baseline) or cue difficulty/reward prediction (AMP instructions, AMP experience). AMP baseline was preceded by a short practice phase of 24 trials.

**Figure 1.**
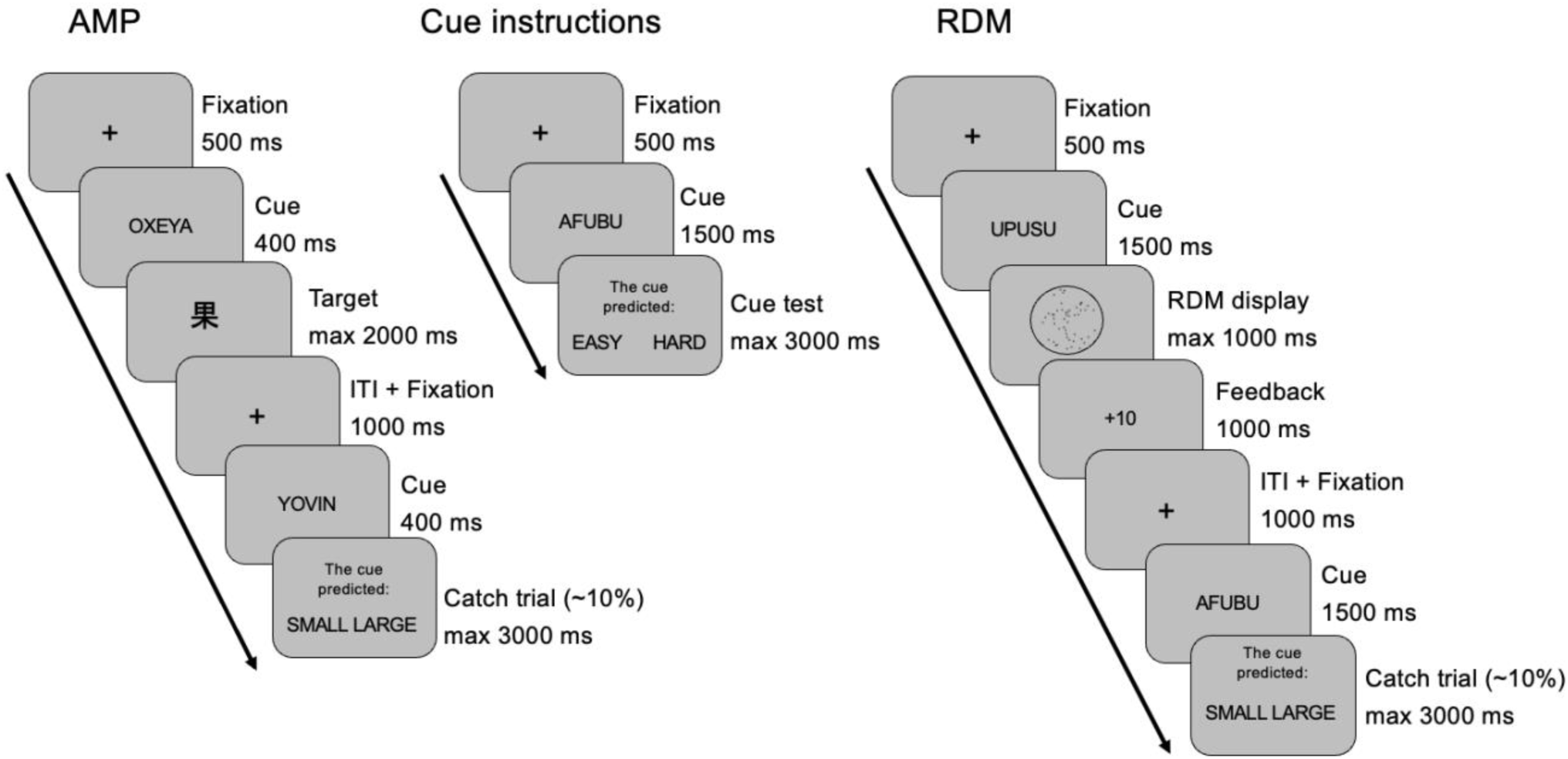
Illustration of experimental procedure. *Note*. Example of trial sequences in different experimental phases. Participants performed a first AMP before learning the meaning of the cues (AMP baseline). In that phase, catch trials tested the identity of the just-displayed cue rather than its meaning. Afterwards, participants learned the meaning of cues through an active learning phase, after which the AMP was administered again (AMP instructions). Finally, participants completed the RDM task and the AMP again (AMP experience).

In the cue instruction phase (Figure 1, central panel), participants learned about the correspondence between the nonword cues and a combination of reward (small or large) and difficulty (easy or hard) levels (counterbalanced between participants). To strengthen the instructions, they were presented across 24 trials with a two-option display probing one dimension of the cue. For example, if the cue anticipated a “hard, small reward” and the reward dimension was probed, they had to select between “small” and “large” options by pressing the C or N key. Each of these trials started with a fixation cross (500 ms), the cue (1500 ms), the two-options display (until response or max 3000 ms) and an interval (500 ms). On a final test of 8 trials, they had to reach 100% accuracy or repeat the test again for a maximum three times after which they could not continue the experiment.

The RDM task (Figure 1, right panel) consisted of 160 trials presented across 4 blocks. Each trial started with a fixation cross (500 ms), cue (1500 ms), random dot motion target (until response or max 1000 ms), feedback (1000 ms), and an interval (500 ms). The target was a dynamic display with a percentage of dots moving coherently towards left or right against a random dot motion background. Participants had to respond to the coherent motion direction by pressing the C or N key respectively. Crucially, the target was preceded by one of the instructed cues predicting difficulty and reward levels of each trial (40 trials each). The difficulty of the task (percentage motion coherence) was adjusted per subject during practice through a staircase procedure converging to 70% accuracy. The resulting coherence was used for the hard level (M=16% coherence, SD=8% coherence), and it was doubled for the easy level. The reward was based on points system (1 point = small reward, 10 points = large reward) which was converted to money at the end of the task (max 7 euro, M=5.82, SD=0.68). Participants received points feedback on each trial (+1, +10, 0) and at the end of each block to keep track of their reward. To promote cue processing, ∼10% of catch trials probed their understanding of either the reward or difficulty level at stake.

The questionnaires administered at the end of the experiment measured individual tendencies to engage and enjoy cognitively demanding tasks (Need For Cognition, NFC, Cacioppo & Petty, 1982), individual tendencies to avoid aversive outcomes and approach goal-oriented outcomes (Behavioral Inhibition System/Behavioral Activation System, BIS/BAS, Carver & White, 1994), individual differences in state and trait anxiety (State-Trait Anxiety Inventory, STAI, Spielberger, 1983), and individual traits of impulsivity and inattention (Adult ADHD Self Report Screen Scale for DSM-5, ASRS-5, Ustun et al., 2017).

### 2.3. Data analysis

#### 2.3.1. Preprocessing

From the initial sample (N=100), 1 participant was excluded due to a server error, for which data was not recorded correctly. We then excluded participants who performed below chance level in response to RDM displays or catch trials in either AMP phase (N=11) or RDM task (N=1). Chance levels were determined by separate binomial tests. We further excluded those who displayed a judgment bias (> 90% negative or positive in either AMP phase as in Vermeylen et al., 2019; N=3). We analysed data from the remaining 84 participants. For RT analyses we excluded errors, post-error trials, trials with responses faster than 200 ms and trials with responses slower or faster than 2.5 SD from the mean RT computed per subject and cue condition. RT were log-transformed before analyses. For the accuracy analyses we excluded trials preceded by an error. For analyses on AMP task, we excluded trials faster than 200 ms.

#### 2.3.2. Statistical models

The main statistical analyses were conducted using Bayesian mixed effects models with brms package (Bürkner, 2021) in R (R Core Team, 2022). For exploratory correlations, Bayes Factors were computed with BayesFactor package.

For the main analysis on AMP data, we ran a model predicting pleasantness of judgements based on phase (3; AMP baseline, AMP instructions, AMP experience), and cue predictions of difficulty (2; easy, hard) and reward (2; small, large). We included random slopes for phase, difficulty and reward and their interaction per participant. To test our hypotheses, we coded our contrasts to compare AMP experience vs AMP baseline (Phase3vs1 contrast) and AMP instructions vs AMP baseline (Phase2vs1 contrast). Pleasantness of judgements was modelled with Bernoulli distribution (logit link).

For the analysis on RDM data, we ran separate models predicting RT and accuracy based on cue predictions of difficulty (2; easy, hard) and reward (2; small, large). We included random slopes for difficulty and reward and their interaction per participant. Log-transformed RTs were modelled with Gaussian distribution (identity link), and accuracy was modelled with Bernoulli distribution (logit link).

For exploratory analyses of individual differences, we included z-transformed scores on questionnaires as continuous predictors in the previous models. Simple correlations between subject means in AMP/RDM performance and subject scores on questionnaires were also run. The relationship between AMP effects and RDM performance was also investigated by the means of correlations.

For all of our analyses we reported the probability of direction (pd) and 95% credible intervals (95% CI) excluding 0 to infer existence and significance of effects, respectively, as suggested by Makowski et al., 2019.

## 3. Results

### 3.1. Performance in cued RDM task

As shown in Figure 2, participants were more accurate (Difficulty, b = -1.36, 95% CI [-1.56, -1.18], pd = 100%) and faster (Difficulty, b = -0.08, 95% CI [-0.09, -0.07], pd = 100%) on easy trials compared to hard trials. More importantly, they were more accurate on large rewards trials compared to small reward trials (Reward, b = 0.13, 95% CI [0.03, 0.23], pd = 99.5%), with no concomitant effect on response times (Reward, b = -0.00, 95% CI [-0.01, 0.01], pd = 50%). The main effects did not interact, neither in accuracy (Difficulty x Reward (b = -0.04, 95% CI [-0.26, 0.16], pd = 65.3%) nor response times (Difficulty x Reward (b = -0.00, 95% CI [-0.01, 0.01], pd = 55%).

**Figure 2.**
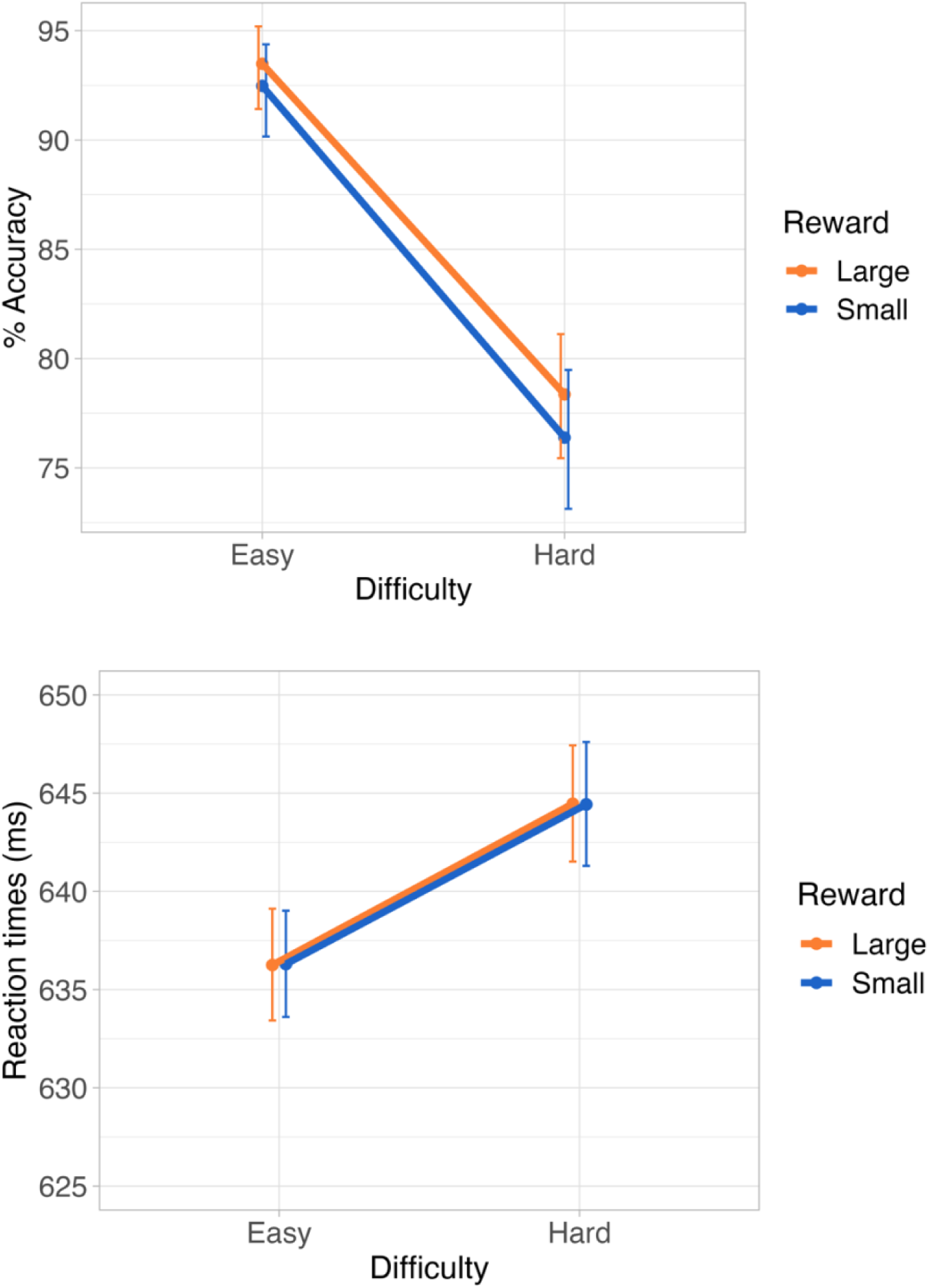
Accuracy and Reaction times results in the RDM task. *Note*. Percentage of correct responses (top) and reaction time measurements (bottom) in response to trials prospecting reward and difficulty cues in the RDM task. Error bars represent 95% credible intervals.

### 3.2. Affective evaluation of cues in AMP

In our main preregistered analysis, we tested whether the AMP effect was already present after being instructed on the meaning of the cues or only after experiencing them in a separate task compared to baseline. Results showed no main effects. However, as illustrated in Figure 3, more negative evaluations were observed for small rewards versus large rewards already after AMP instructions (Phase2vs1 x Reward, b = 0.11, 95% CI [0.03, 0.15], pd = 98%) as well as after AMP experience compared to baseline (Phase3vs1 x Reward, b = 0.08, 95% CI [0.00, 0.16], pd = 100%). These two-way interactions between reward and phase were not modulated by difficulty in either contrast (Phase2vs1 x Reward x Difficulty, b = 0.01, 95% CI [-0.05, 0.06], pd=60%; Phase3vs1 x Reward x Difficulty, b = -0.02, 95% CI [-0.07, 0.03], pd = 75%).

**Figure 3.**
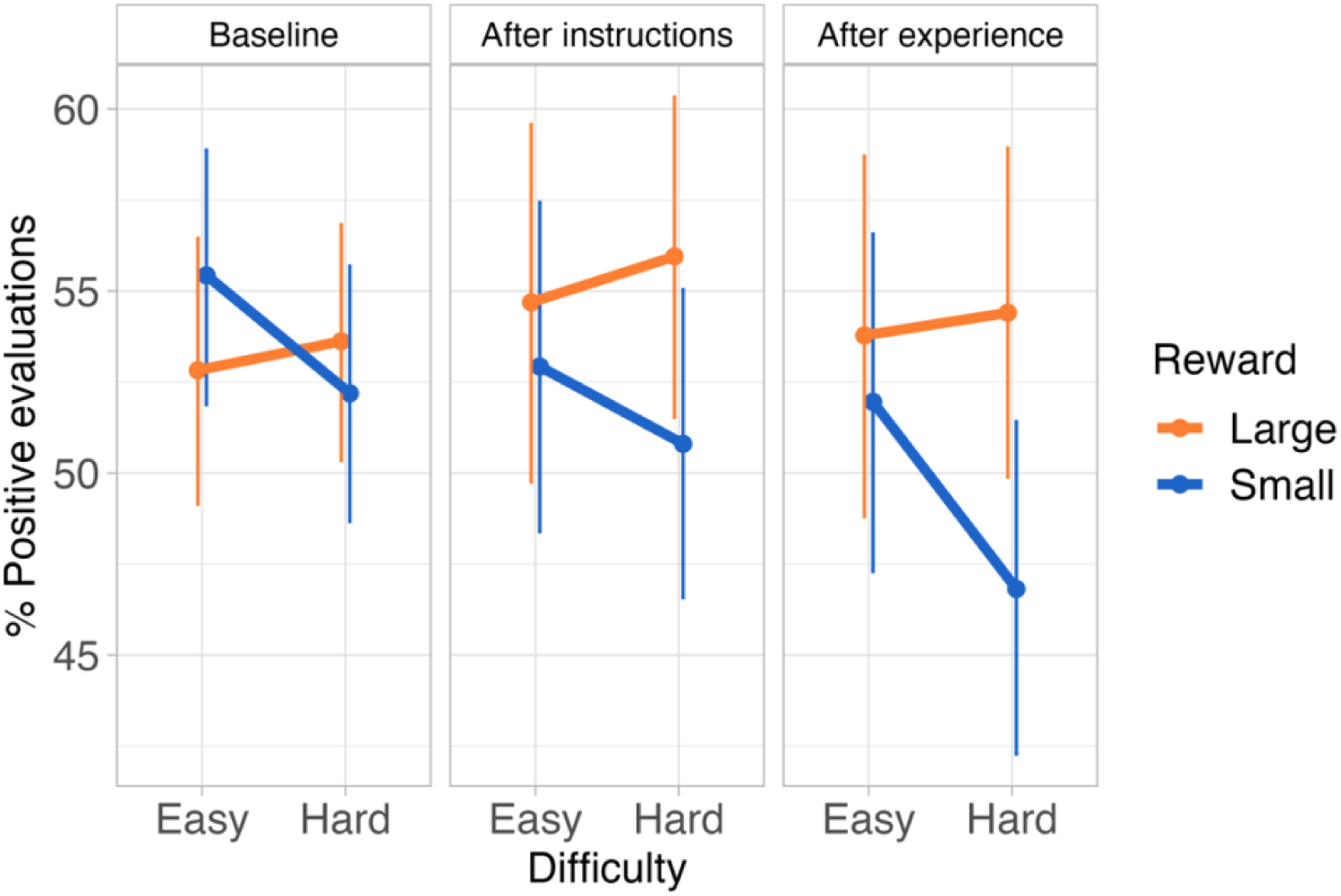
AMP results. *Note*. Percentage of positive evaluations of cues prospecting reward and difficulty levels at baseline (before cue instructions or experience), after instructions of cues, and after experience of cues in the RDM task. Error bars represent 95% credible intervals.

As an exploratory analysis, we wanted to see whether participants significantly changed their evaluations after having experienced the cue in the task compared to instructions only. We subtracted the baseline means per condition from the respective means of the other two phases and ran a Phase (2; AMP instructions vs AMP experience) x Reward (2; small vs large) x Difficulty (2; easy vs hard) Bayesian ANOVA. Results revealed evidence for a reward effect (BF10 = 10) against an intercept only model, independent of phase or difficulty conditions (0.3 <BFs < 1).

### 3.3. Individual differences in the affective evaluation of cues

We hypothesised a positive correlation between the reward effect observed on accuracy in the RDM task and the reward effect in either AMP instruction or AMP experience. However, the analyses did not find any evidence for such correlations (all BF10 < 3).

Likewise, implementing questionnaires scores in models analysing RDM or AMP performance did not show any evidence for an impact of either need for cognition, tendency to approach/avoid goal-oriented outcomes, trait or state anxiety nor traits of impulsivity and inattention on reward effects (all CIs included 0).

Finally, simple correlations between averaged means per subject between questionnaire scores did not show any effect (all BF10 <1)^2^.

## 4. Discussion

In this pre-registered study, we investigated whether large reward expectations are affectively evaluated more positively and mitigate difficulty expectations even in the absence of task experience. To this end, we measured affective evaluations (AMP) of cues prospecting reward and difficulty levels immediately after cue instructions (i.e., without task experience; AMP instructions) as well as after engagement in a random dot motion task (RDM), and we contrasted these conditions to a baseline measurement (AMP baseline), in which the cues were just meaningless nonwords. Compared to baseline, we expected to observe more negative evaluations of small reward cues (possibly even more so on hard trials) not only after task experience (as expected based on Vermeylen et al., 2019; 2022), but, crucially, also immediately after instructions.

Performance in the RDM task confirmed that our difficulty manipulation was successful and that participants paid attention to, and adapted their performance based on cue prospects. In line with previous research investigating the effects of motivation on mental effort investment, participants were more accurate on trials prospecting a large reward compared to a small reward (e.g., Frömer et al., 2021; Kukkonen et al., 2023; Otto & Vassena, 2021). However, reward did not mitigate the effect of task difficulty (no cost/benefit trade-off), nor did it have an impact on reaction times, possibly due to the nature of the task. The RDM task requires participants to accumulate evidence on the global motion direction, and performance is more dependent on the amount of available information (i.e., data-limited) than on the resources allocated to the task (i.e., resource-limited, as can be e.g. an arithmetic task; Norman & Bobrow, 1975). On the one hand, hard conditions in data-limited tasks could have overall been experienced as less aversive and therefore less subjective to the cost-benefit trade-off. On the other hand, evidence accumulation could have counteracted a possible speeding effect after large reward cues. Regardless of whether participants invested more effort when a larger reward was prospected or simply adapted their performance strategy, results in the RDM task confirm that participants used reward cues throughout the task to adjust performance.

Our analyses on their affective evaluations showed that this performance benefit was accompanied by more positive judgements of large reward cues compared to small reward cues. Crucially, compared to baseline measurements, the effect was present both following instructions on the meaning of the cues (AMP instructions) and after participants had experienced the predictive values of cues in the RDM task (AMP experience), with no difference between these two phases. To the best of our knowledge this is the first study to show evaluative effects of reward anticipations in the absence of any task experience. For instance, Van Dessel et al., (2020) observed AMP effects on task switching/repetition cues if participants had a short experience of the task with another pair of cues (Experiment 3, Group 2), whereas the effect disappeared when AMP was measured immediately on the first set of cues, in the absence of any experience (Experiment 3, Group 1). Importantly, the cues in Van Dessel et al., referred to more abstract ‘switching’ and ‘repetition’ experiences, for which a short task practice might be necessary. On the contrary, in our experiment, the cue instructions referred to task difficulty and possible reward gains, which are concepts participants had already learned outside the experimental context. Indeed, a prerequisite for rewards to acquire affective value and motivational salience is to first learn about relationships among stimuli and consequences of actions (Berridge & Robinson, 2003). We are constantly motivated by the prospect of reward – kids get a cookie when we they finish the vegetables, adults get promotions when they work harder – therefore, it is not surprising that a reward cue is appreciated even in the absence of a specific task. Similar to our study, Van Dessel et al., (2017) showed evaluative effects based on instructions for nonwords indicating approach or avoidance behaviours, which represent typical and adaptive action tendencies in the presence of positive or negative stimuli, respectively (Phaf et al., 2014).

Nevertheless, we did not observe a comparable evaluative effect for task difficulty as we did with reward cues. We can only speculate about the cause of this asymmetry, but we assume the relative saliency of both dimensions might differ. Participants were informed from the outset about the possibility of winning a concrete amount of reward (up to 7 extra euros), whereas the instructions regarding task difficulty (i.e., motion coherence) were less tangible and therefore potentially less salient. In addition, previous behavioural and computational modelling results suggest that in general participants prioritise reward over difficulty information when both are simultaneously cued (Vassena et al., 2019).

Taken together, the data suggest that instructed beliefs (on top of previously learned expectations) play a crucial role in the evaluation process preceding mental effort investment. As expected, experiencing the predictive value of cues in a short task did not modulate (i.e., strengthen) the affective evaluation of reward cues. Previous studies measuring the AMP only after task experience showed that affective evaluations can correlate with performance in the main task (Van Dessel et al., 2020; Vermeylen et al., 2019), but not always (Vermeylen et al., 2022). The lack of a consistent correlation, which we also observed in our study, might be attributed to AMP capturing instructed beliefs that remain independent of and unchanged by task experience. Future studies should further investigate whether AMP measurements after task experience (vs. instructions only) are affected and correlate with task performance when more salient manipulations of task difficulty and rewards are used.

Finally, our findings inform existing theories of mental effort investment, such as the EVC, which conceives mental effort as the outcome of an evaluative process weighing costs and potential benefits (Shenhav et al., 2013, 2017). A previous study (Devine et al., 2023) has shown that actual effort investment (as indexed by task performance) is independent from trial-by-trial affective reactions to reward and difficulty cues. Our study contributes by suggesting that affective evaluations and subsequent effort investment may reflect broader instructed expectations related to reward and task difficulty. These evaluations may remain relatively stable and show minimal updates during task performance or even after it. Our suggestion aligns with other studies demonstrating that continuously adapting effort signals on trial-by-trial reward and difficulty information is resource-intensive akin to the costs associated with task switching (Grahek et al., 2022), and that participants rather maintain an effort signal based on block instructions (Kukkonen et al., 2023).

In conclusion, the current study demonstrates for the first time that affective evaluations of reward and difficulty cues (even after task experience) are primarily influenced by instructed expectations related to a task set (e.g., potential rewards, perceived difficulty’s salience) before any actual task exposure. Importantly, these instructed expectations significantly impact the evaluation and subsequent investment of mental effort.

^1^We deviated from our pre-registration and instead of following a Bayesian sequential sampling design with an ultimate stopping rule at 100 participants, for practical constraints, we directly collected data from 100 participants to maximise the power of our analyses.

^2^We observed a positive correlation between the AMP effect (after instructions as well as after experience) and scores on NFC questionnaire (BF=4.81 and BF=8.96, respectively). However, the correlation disappeared (BF=0.61, and BF=0.33, respectively) when outliers were accounted for (+-2.5 SD; N=3).

